# Metabolomic fingerprints of PAH exposure - identifying toxicological biomarkers in dynamically cultured 3D cell spheroids

**DOI:** 10.1101/2025.07.10.663939

**Authors:** Anja Bosnjakovic, Thomas O. Eichmann, Alja Stern, Morten Arendt Rasmussen, Mario Lovric, Bojana Zegura

## Abstract

Environmental exposure to polycyclic aromatic hydrocarbons (PAHs) causes metabolic dysfunction, but reliable biomarkers are still needed to assess human health effects. This study used 21-day matured human HepG2 spheroids, a metabolically competent three-dimensional (3D) liver model, to assess metabolic responses to graded, non-cytotoxic concentrations of benzo[a]pyrene (BaP) and benzo[b]fluoranthene (BBF) after 24- and 96-h exposure. Untargeted liquid chromatography-mass spectrometry (LC–MS) metabolomics, combined with multivariate and network analyses, identified compound- and time-specific metabolic signatures. At 24 hours, no metabolites showed significant changes. In contrast, at 96 hours, both PAHs consistently altered seven robust metabolites linked to polyamine metabolism, membrane dynamics, mitochondrial energy, and DNA-repair pathways. Network analysis showed BBF caused broader and more connected changes than BaP, indicating distinct toxicodynamics. These findings underscore the importance of extended exposure in revealing metabolic disruption and support a set of candidate biomarkers for future low-dose studies and improved risk assessment of airborne toxicants.

## Introduction

Polycyclic aromatic hydrocarbons (PAHs) are ubiquitous environmental pollutants generated primarily through the incomplete combustion of organic materials (Choi et al., 2010). Human exposure to PAHs occurs via multiple pathways, notably through inhalation of contaminated ambient and indoor air, ingestion of tainted water and food, and dermal contact with polluted soils and dust (Mallah et al., 2022). In urban and industrial environments, PAHs are often adsorbed onto particulate matter, thereby posing significant health risks. Chronic exposure to these compounds has been implicated in ethology of various malignancies, including cancers (Jakovljević et al., 2025; Štampar et al., 2021), as well as cardiovascular diseases (Mallah et al., 2021). The toxicological impact of PAHs is largely attributed to their metabolic activation by xenobiotic-metabolizing enzymes, leading to the formation of reactive intermediates capable of inducing oxidative stress (Ryu and Hong, 2024) and forming DNA adducts (Ewa and Danuta, 2017). These interactions can result in genomic instability (Bai et al., 2017; Kazensky et al., 2024), thereby enhancing the risk of mutagenesis and carcinogenesis. Despite the extensive studies on the toxic effects of PAHs, a critical gap remains in the identification and validation of reliable biomarkers for exposure and early biological effects, particularly at lower concentrations. Traditional risk assessments such as the frequently used incremental lifetime cancer risk (ILCR) (Jakovljević et al., 2020; Lovrić et al., 2024c) have predominantly relied on data from the 1990s (Nisbet and LaGoy, 1992), underscoring the urgent need for updated approaches. Although genotoxicity assays and DNA damage evaluations have provided valuable information on PAH-induced adverse effects, a comprehensive systems-level approach is crucial to fully capture the complex biochemical perturbations triggered by PAH exposure. Metabolomics, the large-scale study of metabolites within biological systems, offers a powerful platform identifying novel biomarkers and elucidating metabolic alterations associated with PAH exposure (Chen et al., 2017; Gao et al., 2018), as well as other xenobiotics such as PFAS (Guo et al., 2022; India-Aldana et al., 2023; Lovrić et al., 2024b) and phthalates (Deng et al., 2024; Fang et al., 2024). By providing a comprehensive snapshot of metabolic responses, metabolomics can enhance our understanding of PAH toxicity mechanisms and contribute to more accurate exposure assessments, even at the low concentrations commonly encountered in indoor environments (Lovrić et al., 2024a). Among the numerous PAH compounds, benzo[a]pyrene (BaP) and benzo[b]fluoranthene (BBF) (IARC, n.d.) stand out due to their high toxicological relevance and widespread presence in the environment (Authority (EFSA), 2008). BaP is a well-characterized genotoxic and carcinogenic agent, classified as a Group 1 carcinogen by the International Agency for Research on Cancer (IARC, n.d.). It serves as a reference compound in many regulatory frameworks for assessing PAH-related risks, owing to its extensively studied metabolic activation pathways and its ability to form DNA adducts that initiate mutagenesis. In contrast, BBF, classified by IARC as possibly carcinogenic to humans (Group 2B) (IARC, n.d.), remains considerably less studied than BaP, despite being one of the 16 polycyclic aromatic hydrocarbons designated as priority pollutants by the U.S. Environmental Protection Agency due to their toxicity and environmental persistence (US EPA, 2013). Nevertheless, it is increasingly recognized for its significant contribution to PAH-associated toxicity, particularly in urban air. Experimental studies have shown that BBF can induce oxidative stress (Bao et al., 2023), mutations (Schuster et al., 2024), DNA strand breaks, DNA adducts (Topinka et al., 2008), and chromosomal alterations (Schuster et al., 2024), indicating its genotoxic potential. At higher doses, BBF exhibits a strong mutagenic response particularly in the liver, an organ with high metabolic capacity, primarily inducing C:G > A:T transversions, followed by C:G > T:A and C:G > G:C substitutions, suggesting preferential targeting of guanine and cytosine bases. Its mutation spectrum correlates with tobacco-related mutational signatures, reinforcing BBF’s role in the carcinogenicity of combustion-derived PAHs and showing mutagenic effects comparable to those of benzo[a]pyrene, a well-characterized reference PAH (Schuster et al., 2024). These findings underline the need for further investigation into BBF’s biological effects, as the available toxicological data remains less comprehensive than those available for BaP. A deeper understanding of BBF’s mechanisms of action will be crucial for refining hazard assessments and understanding its relative toxicity in comparison to better-characterized PAHs like BaP.

In this study, we employ high-resolution metabolomic profiling of PAH-exposed 3D cell models to identify specific metabolic signatures indicative of PAH exposure using liquid chromatography-mass spectrometry (LC-MS/MS) analysis. By integrating comprehensive statistical methods and machine learning approaches, we aim to uncover robust biomarkers and gain deeper insights into the biochemical pathways perturbed by PAHs. Given the limitations of conventional two-dimensional (2D) cultures in replicating physiological responses, we apply a three-dimensional (3D) spheroid model generated in a clinostat-based system (CelVivo). This advanced, state-of-the-art technology enables the formation and long-term cultivation of uniform and functional spheroids that closely mimic *in vivo* tissue architecture, cellular heterogeneity, and particularly metabolic activity found in human liver tissues (Wrzesinski and Fey, 2018). Notably, in 3D cultures the metabolic competence of immortalized liver cell lines is significantly enhanced compared to 2D monolayer cultures, overcoming the limitations of compromised xenobiotic metabolism often observed in 2D models. This is particularly important when assessing the adverse effects of xenobiotics like polycyclic aromatic hydrocarbons, which require metabolic activation to exert their toxicity. Moreover, the ability to maintain the spheroids under dynamic, controlled conditions further allows for prolonged exposures, making this system particularly suitable for studying chronic effects of environmental stressors (Štampar et al., 2021; Štampar and Žegura, 2024). By using this innovative system, we improve the physiological relevance and translational potential of our *in vitro* toxicological assessments, offering a closer approximation of real-world human exposure scenarios.

This research seeks to advance the identification of reliable biomarkers for human exposure to PAHs, thereby contributing to improved future risk assessment strategies and the refinement of regulatory policies addressing PAH-related health risks. To the best of our knowledge, this is the first metabolomics-driven investigation to systematically characterize PAH-induced metabolic perturbations using 3D hepatic spheroid models. These findings have the potential to deepen our understanding of PAH toxicity mechanisms and support the development of more accurate biomarkers for future research and risk assessment.

## 2. Materials &Methods

### 2.1. Cell culture

The immortalized human hepatocellular carcinoma cell line HepG2 was purchased from the American Type Culture Collection (ATCC-HB-8065™, Manassas, VA, USA). The cells express wild-type TP53 and retain several xenobiotic-metabolizing enzyme activities; therefore, they are widely used as an *in vitro* model for toxicological studies. Cells were cultured in Minimum Essential Medium (MEM, 10370–047; Gibco, Paisley, Scotland), supplemented with 10 % foetal bovine serum (FBS; Gibco), 1 mM sodium pyruvate, 2 mM L-glutamine, and 100 IU/ml penicillin/streptomycin (all from Sigma-Aldrich, St. Louis, MO, USA). Cultures were maintained at 37 °C in a humidified atmosphere with 5 % CO₂. HepG2 cells between passages 3 and 10 were used for the experiments at approximately 70 % confluency. Mycoplasma contamination was routinely monitored using the MycoAlert™ Mycoplasma Detection Kit (Lonza, Walkersville, MD, USA).

### 2.2. Development of 3D spheroids and culture conditions

For experimental purposes, 3D cell models (spheroids) were generated by pre-forming cell aggregates using AggreWell™ 400 µm 24-well plates (STEMCELL technologies, Vancouver, Canada) following the manufacturer’s instructions. Cells were seeded at a density of 200.000 cells/well, yielding approximately 170 cells per microwell, in complete growth medium, followed by centrifugation at 100 g for 5 min. The plates with spheroids were incubated overnight at 37 °C in a humidified atmosphere containing 5 % CO_2_ to allow for cell self-aggregation. Spheroids were subsequently cultured under dynamic conditions using the ClinoStar 2™ rotating bioreactor system (CelVivo ApS, Odense, Denmark). To prepare the single-use ClinoReactors^TM^ (CelVivo ApS, Odense, Denmark), sterile water was added to the humidification beads located in the humidity chamber, allowing them to hydrate at room temperature for 4 hours. Once hydrated, 7 mL of HepG2 growth medium was added to each ClinoReactor^TM^. The units were then placed in ClinoStar 2^TM^ system and left to equilibrate overnight at 37 °C in a 5 % CO₂ atmosphere, rotating at a speed of 12 rpm. On the following day (Day 0), cell aggregates were gently detached from the AggreWell™ plates by pipetting, inspected under a light microscope to determine their quality, and transferred (approximately 1200 cell aggregates prepared in a single well per ClinoReactor^TM^) into the pre-equilibrated ClinoReactor^TM^. The bioreactors were then filled with growth medium to a final volume of 10 mL and placed into the ClinoStar 2^TM^ system, where they were maintained at 37 °C in a 5 % CO_2_ atmosphere under continuous rotation (initial rotation speed of 10 rpm). To support optimal spheroid growth, 90 % of the culture medium was refreshed on a 48/48/72-hour schedule. The rotation speed of ClinoReactors^TM^ was adjusted progressively to accommodate spheroid growth. During each media change, a “gardening” step was performed to remove spheroids that were irregular in size or shape, promoting greater uniformity within the culture. The spheroids were transferred to new equilibrated ClinoReactor^TM^ units at day 10 to ensure optimal growth conditions, prevent the accumulation of cellular debris, and maintain the integrity of the experimental environment. HepG2 spheroids were cultured over a 21-day period before being exposed to the studied PAH compounds.

### 2.3. Exposure of HepG2 spheroids to BaP and BBF

After 21 days of dynamic culture, mature spheroids were transferred to U-bottom 96-well low attachment plates (1 spheroid/ well; Falcon, Corning Corporation, New York, USA) and exposed to the selected PAH compounds under static conditions for 24 and 96 h. The spheroids were treated with two concentrations of benzo[a]pyrene (BaP) and benzo[b]fluoranthene (BBF) (Sigma-Aldrich, St. Louis, MO, USA): 20 µM and 2 µM for the 24-hour exposure, and 10 µM and 1 µM for the 96-hour exposure. Corresponding solvent controls were included: 0.2 and 0.05 % for 24h and 0.1 and 0.025 % for 96h, for BaP and BBF, respectively.

### 2.4. Cell viability in spheroids following BaP and BBF – A treatment TP assay

To confirm that the tested concentrations of BaP and BBF did not induce cytotoxic effects, cell viability was assessed using the CellTiter-Glo® 2.0 Cell Viability Assay (Promega, Madison, Wi, USA). This luminometric assay quantifies ATP levels that correlate with the number of viable cells. The assay was conducted following the manufacturer’s protocol, with slight modifications. Briefly, at least three spheroids per experimental point (one spheroid per well) in 50 µl of cell culture media were placed into a white-opaque 96-well plate (Corning, Corning, NY, USA), and 50 µl of the assay reagent was added to each well. The content of each well was thoroughly mixed by pipetting, and the reaction was incubated at RT for 20 min. Luminescence was then measured using a luminometer (Sinergy HTX, BioTek, Winooski, VT, USA). Raw data were normalized to the mean values of the corresponding solvent controls. GraphPad Prism v10 software (GraphPad Software, San Diego, CA, USA) was used to identify significant differences in cell viability between PAH-exposed spheroids and corresponding solvent controls, a One-Way Analysis of Variance (ANOVA) followed by Holm-Šídák’s multiple comparison test was performed. The experiments were repeated three times independently (*p < 0.05).

### 2.5. Sample collection for metabolomic analysis

For metabolomic analysis, 40 spheroids per experimental point were collected into 2 ml Safe-Lock Eppendorf tubes (Eppendorf, Hamburg, Germany). The cell media was discarded, and spheroids were washed with 1x PBS (1 ml). After the washing step, the PBS was removed entirely from the tubes. The spheroids were then immediately snap-frozen by submerging the tubes in liquid nitrogen for 20 seconds. Frozen samples were stored at –80 °C until metabolomic analysis. The experiments were repeated three times independently. A schematic overview of the experimental design is presented in Figure 1.

**Figure 1.**
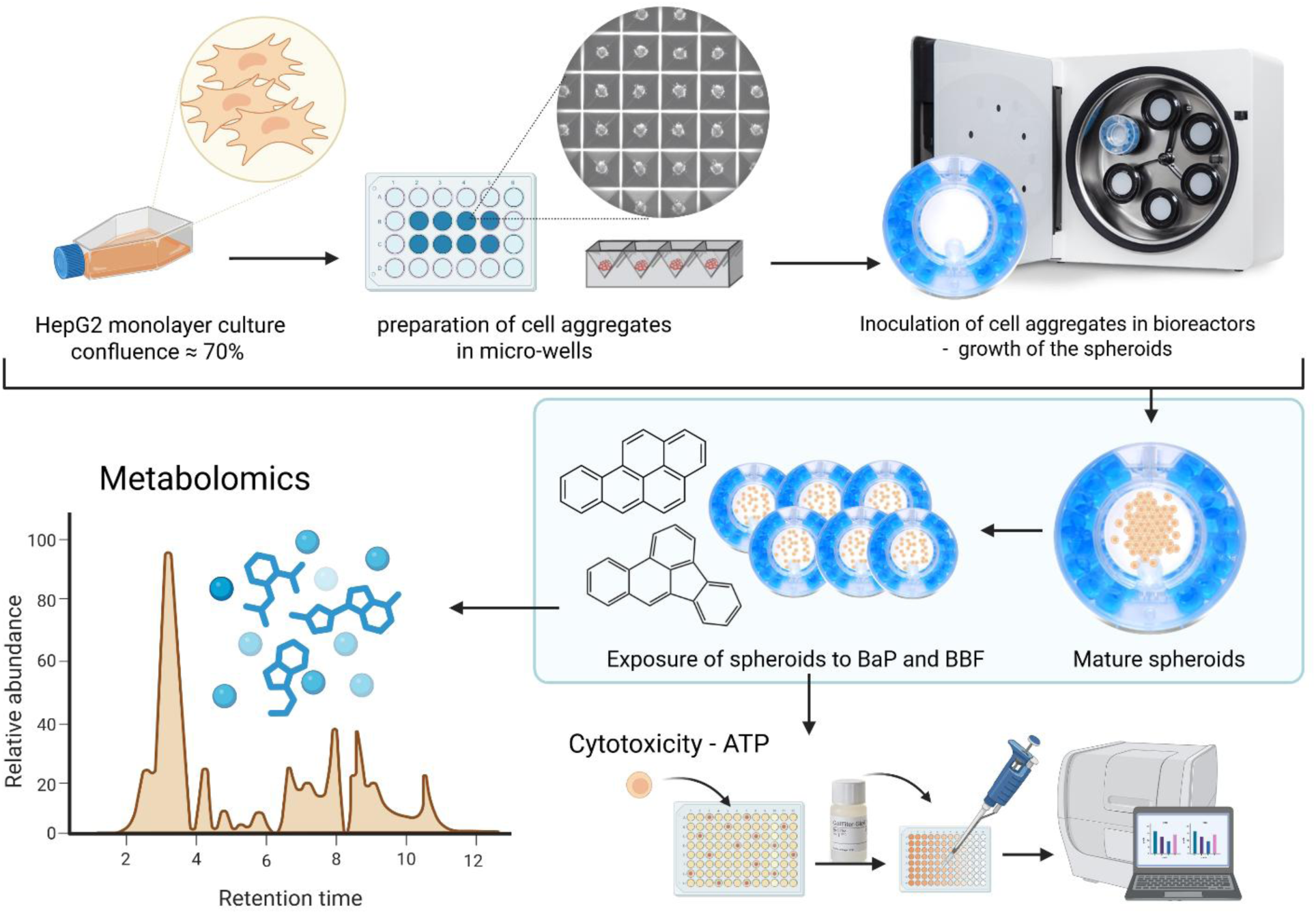
A schematic overview of the experimental design.

### 2.6. Metabolite extraction

Spheroid samples (pooled per condition) were transferred to 2 ml Safe-Lock polypropylene tubes on 4 °C, mixed with 400 µl MeOH (-80 °C) and 100 µl ddH2O containing internal standards (IS, 4 °C; alanine-¹³C₂, serine-¹³C₂, cholic acid-d₄, taurine-d₄, creatinine-d₃, tryptophan-d₅, lysine-d₂, hypoxanthine-d₂, deoxyglucose-6-phosphate, palmitic acid-¹³C₄ all purchased from Sigma-Aldrich or Biomol). Samples were then sonicated using a Bioruptor Pico (30 min, 4 °C, 30 sec ON/30 sec OFF, frequency high; Diagenode). Following sonication, 300 µl ddH2O and 900 µl methyl tert-butyl ether (MTBE) were added to each sample, and the mixture was incubated under constant shaking (1000 rpm, 15 min, 4 °C). After centrifugation (13,000 rpm, 10 min, 4 °C), 800 µl of the upper phase was removed and replaced by 800 µl of artificial upper phase (prepared from MTBE/MeOH/ddH2O in a 9/4/4, v/v/v ratio). After a second incubation and centrifugation, as described above, the complete upper phase was discarded, and 800 µl of the lower phase was collected and dried using a SpeedVac (Thermo Fisher Scientific). The metabolites were reconstituted in 135 µl of 70 % acetonitrile containing 0.6 mM medronic acid and subsequently used for LC-MS analysis. Protein precipitates were dried, solubilized in 250 µl of 0.3 N NaOH (55 °C, ∼ 4 h), and the protein content was quantified using Pierce™ BCA reagent (Thermo Fisher Scientific) according to the manufacturer’s guidelines.

### 2.7 Metabolome analysis

Aliquots of all sample extracts were pooled to create a quality control (QC), which was analysed at the start and end of the sequence as well as after each 5^th^ sample injection. Two blank extraction samples (prepared without any biological sample, for background control) were injected before and after all sample injections. An in-house reference compound mix (for level 1 identification) was injected at the start of the sequence. Each blank injection was followed by two QC injections for column re-equilibration. Chromatographic separation was performed using a Vanquish UHPLC+ system (Thermo Fisher Scientific) equipped with an ACQUITY UPLC BEH Amide column (2.1 × 150 mm, 1.7 µm; Waters), using an 18 min gradient (negative mode, 400 µl/min) from 97 % solvent A (ACN/ddH2O, 95/5, v/v; 10 mM NH4FA, 10 mM NH3) to 65 % solvent B (ddH2O/ACN, 95/5, v/v; 20 mM NH4FA, 20 mM NH3) or a 23 min gradient (positive mode, 400 µl/min) from 99 % solvent A (ACN/ddH2O, 90/10, v/v; 10 mM NH4FA, 0.1 % FA) to 50 % B (ddH2O, 10 mM NH4FA, 0.1 % FA). The column compartment was kept at 40 °C and 30 °C, respectively. Metabolite detection was performed on a QExactive Focus mass spectrometer (Thermo Fisher Scientific) equipped with a heated electrospray ionization (HESI II) source, operating in both negative and positive data-dependent acquisition (DDA) modes. Metabolites were identified either level 1 via accurate m/z of the [M-H]^-^ or [M+H]^+^ ion (<5ppm) and comparison of the retention time (rt) and MS2 spectra to synthetical reference compounds or level 2 without retention time reference. All QC peaks were manually inspected using Freestyle (1.8 SP2), and peak extraction was performed in Skyline (24.1.0.199). Only metabolites with <25 % peak area variation in QC samples were used for further processing. Peak areas were blank subtracted and normalized to IS (rt-range clustered) and protein concentration. Metabolite levels were expressed in arbitrary units (AU), calculated as the analyte-to-IS ratio normalized per µg of protein.

### 2.8. Statistical analysis and machine learning

To identify metabolites significantly altered by chemical exposure, we applied a conservative and structured statistical workflow. Initially, metabolites were analysed for correlation, with one removed per pair if the Spearman correlation would be above 90 %. Concentration values were then log10-transformed. Samples were grouped according to experimental condition, defined by chemical identity (e.g., BaP or BBF), concentration (low or high), and exposure duration (24 h or 96 h), referred to as Pools in Table 1. Within each experimental context, the metabolite relative concentrations were fed to analysis of variance (ANOVA) to assess whether metabolite levels differed significantly across the three exposure conditions (low concentration, high concentration, and corresponding vehicle control). To control for false positives arising from multiple testing, p-values were adjusted using the Benjamini–Hochberg false discovery rate procedure at alpha 0.05. Metabolites considered to be robust were those which were either significant only at the high concentration, or both concentrations in the same direction (down-or upregulated) comparing to control. To investigate the co-regulation structure among metabolites, we constructed a correlation-based network. Pairwise Spearman correlation coefficients were computed between all metabolites to assess monotonic relationships while mitigating sensitivity to outliers. A correlation threshold of |r| > 0.6 was applied to retain only strong associations; edges were drawn between metabolite pairs exceeding this threshold, and the absolute correlation coefficient was presented as an edge. The resulting undirected, weighted network was constructed using the NetworkX Python library (Hagberg et al., 2008). Nodes represented individual metabolites, and edges represented high-confidence correlations. Community structure within the network was identified using the greedy modularity maximisation algorithm (Clauset et al., 2004), which partitions the graph into non-overlapping clusters (communities) by maximising the density of within-group connections relative to between-group links. To facilitate interpretation, the network was visualised. To further explore the global metabolic differences induced by prolonged exposure to PAHs as a more explorative approach, we performed principal component analysis (PCA) after z-score normalisation of metabolite abundance data. PCA is a dimensionality reduction technique that transforms complex, high-dimensional data into principal components (Wold et al., 1987) capturing the greatest variance in the dataset. By projecting data into this new coordinate space, PCA enables visualisation of patterns, group separation, and the influence of individual variables in a much lower dimensionality, hence understandable to the human eye. The dimensionality reduction was performed on the metabolites following 96-hour treatments with BaP or BBF (1 µM and 10 µM), alongside the matched vehicle controls (VC). We further performed enrichment analysis using the Over Representation Analysis module in MetaboAnalyst 6.0 (Pang et al., 2024), based on the results of our statistical analysis where we identified the most robust upregulated or downregulated metabolites.

**Table 1.** The experimental design for HepG2 spheroid exposure to benzo[a]pyrene (BaP) and benzo[b]fluoranthene (BBF). Spheroids were exposed to each PAH at two concentrations (high and low) and two time points (24 h and 96 h). Each chemical exposure group included a corresponding vehicle control (VC; DMSO at matched concentration). The matrix summarizes the distribution of experiments (Exp. 1.1 – 4.3), including the specific chemical concentration exposure time, and the pooling strategy used for statistical analysis.

|  | Vehicle Control Conc.<br>(DMSO) | Concentration (µM) |  | Time (h) | Experiment<br>(Pools) |
| --- | --- | --- | --- | --- | --- |
|  |  | High | Low |  |  |
| **BaP** | 0.2 % | 20 | 2 | 24 | 1.1, 1.2, 1.3 |
|  | 0.1 % | 10 | 1 | 96 | 2.1, 2.2, 2.3 |
| **BBF** | 0.05 % | 20 | 2 | 24 | 3.1, 3.2, 3.3 |
|  | 0.025 % | 10 | 1 | 96 | 4.1, 4.2, 4.3 |

## 3. Results

### 3.1 Toxicity experiments and viability assessments

The cytotoxicity of benzo[a]pyrene (BaP) and benzo[b]fluoranthene (BBF) was evaluated in HepG2 spheroids using the CellTiter-Glo® 2.0 Cell Viability Assay after 24 and 96 hours of exposure to 2 and 20 µM, and 1 and 10 µM, respectively (Figure 2). After 24 hours, cell viability remained above 90 % for all conditions except BBF at 20 µM, which reduced viability to approximately 80 %. At 96 hours, no cytotoxic effects were observed, and viability levels were comparable to or slightly higher than the controls. According to the predefined threshold of 75 % viability for further analysis, all tested conditions met the criterion, indicating acceptable cytotoxicity in this 3D liver model. Therefore, these concentrations were selected for subsequent metabolomics experiments.

**Figure 2:**
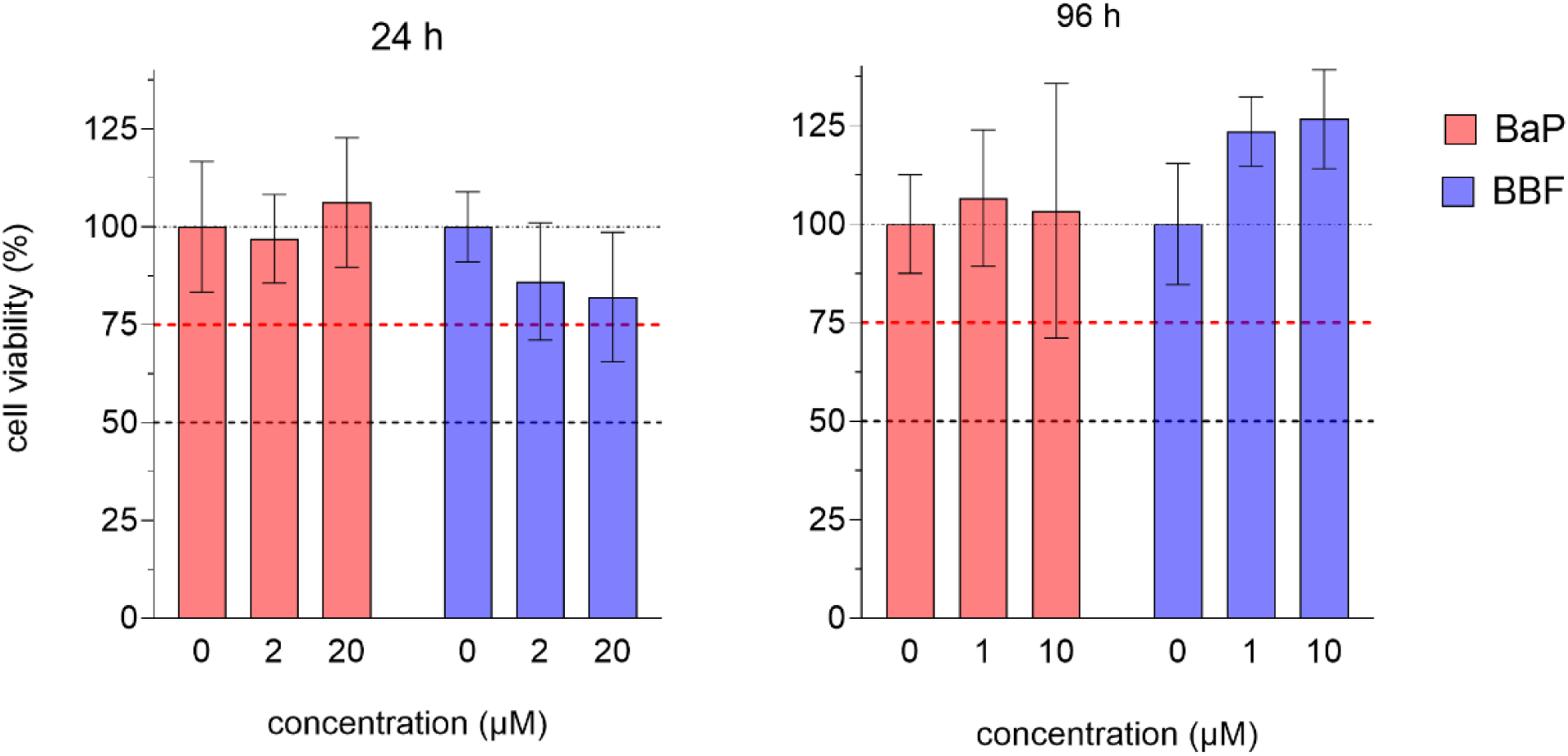
Cytotoxicity of BaP and BBP in HepG2 spheroids after 24 (left) and 96 hours (right) of exposure. Cell viability was evaluated by measuring the ATP content in the spheroids, using the CellTiter-Glo® 2.0 Assay. Cell viability is expressed as the percentage of the solvent control (0; different concentrations of DMSO, corresponding to the concentration present in the samples - for 24 hours: 0.1 % (BaP), 0.05 % (BBF) and for 96 hours: 0.05 % (BaP), 0.025 % (BBF). The dashed lines represent the 100, 75 and, 50 % viability thresholds.

### 3.2 Pattern analysis and metabolite extrapolation

To explore differences in metabolic responses across treatment groups, we performed Principal Component Analysis (PCA) on the metabolite expression data from the 96-hour treatment samples. The resulting biplot (Figure 3) illustrates how samples cluster based on their metabolic profiles following exposure to BaP and BBF at both high and low concentrations, as well as their respective vehicle controls.

**Figure 3:**
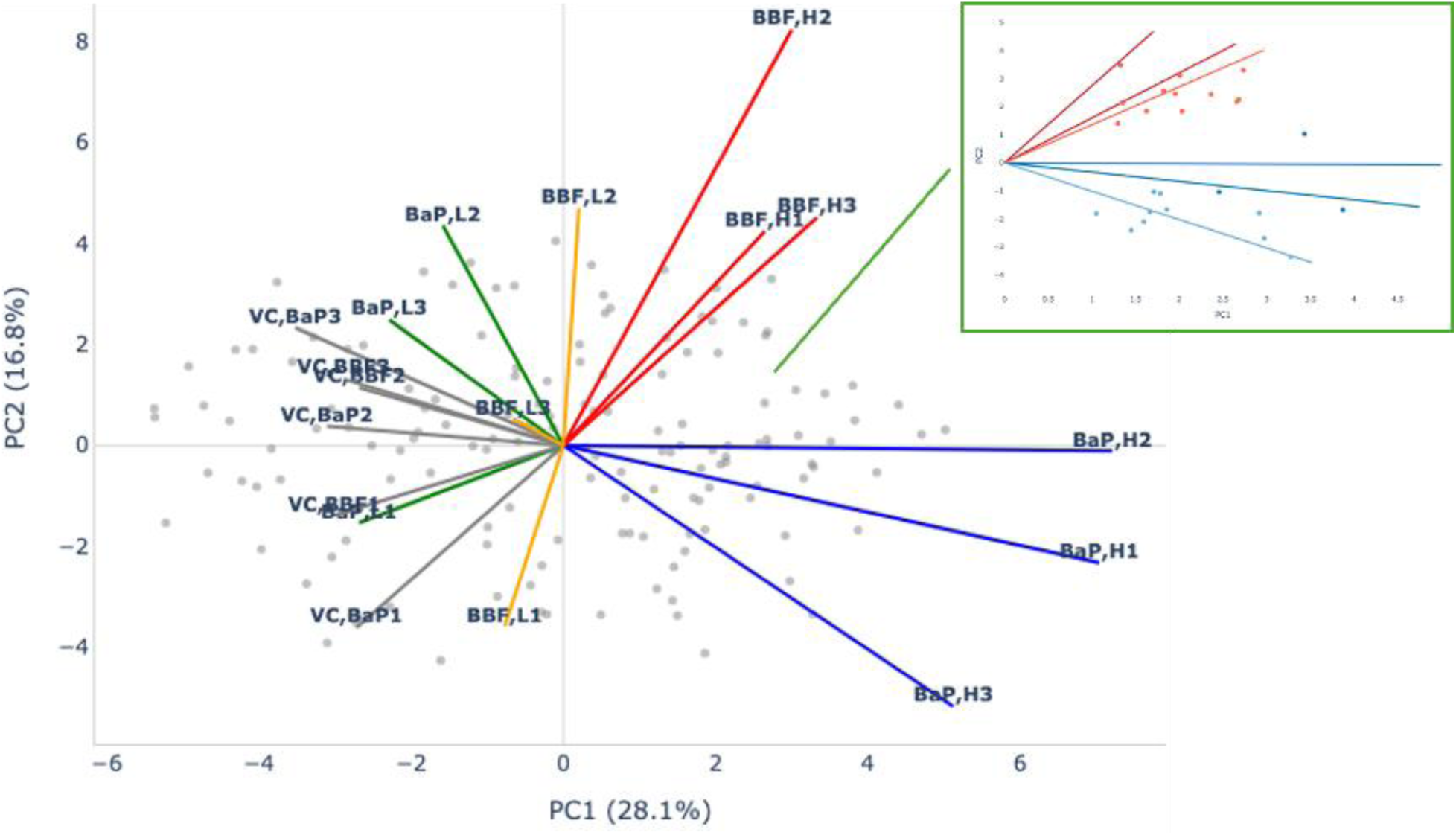
PCA biplot where BaP- and BBF-treated samples cluster separately, with different metabolite groups driving the differences between these treatments. The lines in the biplot represent the loadings, indicating both the direction and relative magnitude of each metabolite’s influence on the principal components.

The high 96-h set was chosen to emphasize the stronger effects in a complex high-dimensional space. The x-axis (PC1) and y-axis (PC2) represent the two main directions of variation in the data. Each point represents a single metabolite, and each coloured vector shows the average response for a specific exposure condition (BaP or BBF at high or low concentration, or control). The direction of a vector indicates the overall type of metabolic change linked to that exposure, while the length shows how strong that effect is. Longer vectors mean a bigger shift in metabolism. The insert in the top right zooms in on the area where the most strongly affected metabolites are found. Metabolites aligned with the BBF high-concentration vector are shown in red, and those aligned with the BaP high-concentration vector are shown in blue. High-concentration BBF samples (red vectors, labelled BBF, H1–3) cluster together and point toward the upper right (positive PC1 and PC2), showing a strong and consistent metabolic response. High-concentration BaP samples (blue vectors, labelled BaP, H1–3) also form a tight cluster, but their vectors point toward the lower right (positive PC1, negative PC2), indicating a metabolic response that is strong but different from BBF. Vehicle controls (grey vectors) point to the left (negative PC1) and stay close to the center on PC2, reflecting minimal changes in metabolism, as this is the non-exposed group. Low-concentration exposures, green for BaP, L1–3 and orange for BBF, L1–3, are also located on the left side of the plot (negative PC1), but their directions along PC2 vary, suggesting weaker or more variable effects at these doses. Overall, PC1 separates samples based on how strong the metabolic response is (higher PC1 values reflect stronger effects), while PC2 distinguishes between the type of response—higher values for BBF and lower for BaP.).

To identify which metabolites were most strongly linked to the high-dose effects of BBF and BaP, we looked for those pointing in a similar direction as the high-dose BBF or BaP vectors in the PCA plot. We measured this using cosine similarity. A value close to 1 means the metabolite’s response closely matches the overall treatment pattern. We selected metabolites with a cosine similarity of at least 0.95 to either the BBF_H or BaP_H (average) vector. We then filtered further, keeping only those located in the upper-right part of the PCA plot (where both PC1 and PC2 are greater than 1), as these contribute the most to separating the treatment groups. This selection resulted in 25 metabolites, shown in the insert of Figure 3. Metabolites aligned with the BBF high-dose response are shown in red, while those aligned with the BaP high-dose response are shown in blue. Of the 25, 12 were most similar to the BBF_H profile, and 13 to the BaP_H profile (see Table 2). These metabolites were used in the network analysis, while others were considered secondary based on further robustness checks.

**Table 2.** Metabolites selected from the PCA given the criterion |PC1| > 1 and |PC2| > 1 from Figure 3 showing high cosine similarity to BaP and BBF high-concentration exposure.

| Metabolite | Aligned | PC1 | PC2 | Metabolite | Aligned | PC1 | PC2 |
| --- | --- | --- | --- | --- | --- | --- | --- |
| N-acetylputrescine | BaP,H | 3.87 | -1.68 | uracile | BBF,H | 2.68 | 2.25 |
| glutamyleucine | BaP,H | 2.46 | -1.04 | pyridoxal | BBF,H | 2.73 | 3.30 |
| docosapentanoic acid | BaP,H | 3.43 | 1.03 | uridine monophosphate | BBF,H | 1.33 | 3.48 |
| stearic acid | BaP,H | 1.66 | -1.75 | phosphate | BBF,H | 1.30 | 1.41 |
| linolenic acid | BaP,H | 1.86 | -1.67 | CDP-ethanolamine | BBF,H | 1.35 | 2.14 |
| myristic acid | BaP,H | 1.78 | -1.09 | UDP-glucose | BBF,H | 2.66 | 2.18 |
| panthenol | BaP,H | 1.45 | -2.41 | cytidine monophosphate | BBF,H | 2.37 | 2.44 |
| citrate | BaP,H | 2.91 | -1.78 | choline | BBF,H | 2.01 | 3.12 |
| fructose-6-phosphate | BaP,H | 3.27 | -3.35 | thiazolidine-4-carboxylic acid | BBF,H | 1.63 | 1.85 |
| oleoylglycerol | BaP,H | 1.05 | -1.81 | acetylcarnitine (C2) | BBF,H | 1.96 | 2.46 |
| urocanic acid cis | BaP,H | 1.60 | -2.09 | glutathione | BBF,H | 2.03 | 1.84 |
| prolylisoleucine | BaP,H | 2.97 | -2.69 | CDP-choline | BBF,H | 1.82 | 2.55 |
| glycine | BaP,H | 1.71 | -1.04 |  |  |  |  |

### 3.4 Statistical Outcomes

The initial step was to explore the ANOVA tested metabolites across all treatment groups. Of the total of 148 metabolites analysed, 20 were differentially in abundance within the BBF | 96h group, 2 in the BaP | 24h group, and 24 in the BaP | 96h group (FDR-adjusted p-value < 0.05), see Figure 4.

**Figure 4.**
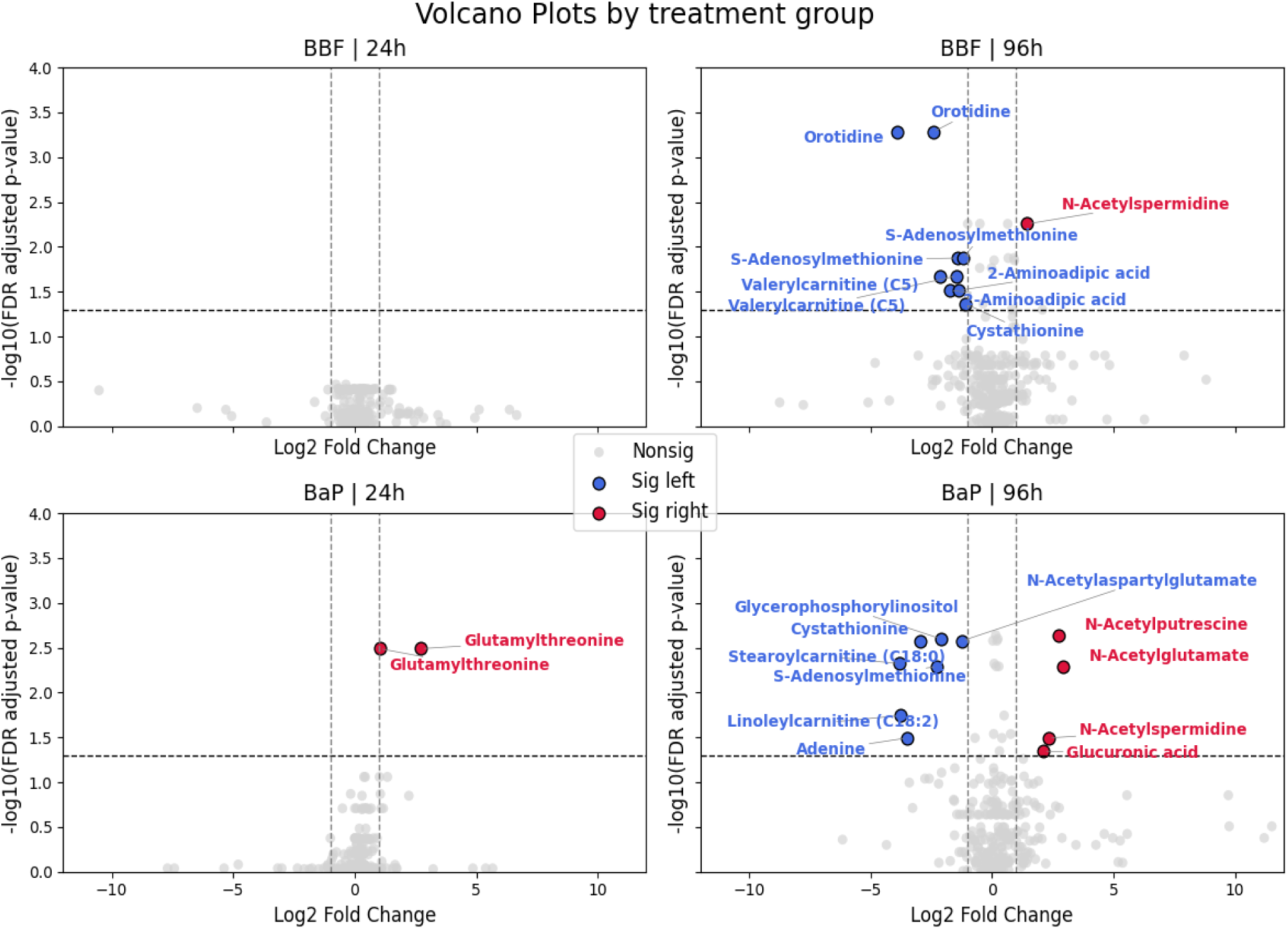
Volcano plots of metabolite changes by group. Volcano plots are showing significant metabolite changes after BaP and BBF exposure at 24 h and 96 h. Red and blue indicate increased and decreased abundance, respectively. Some metabolites (e.g., orotidine) appear more than once due to differential detection across both low and high exposure concentrations.

In the Figure 4, each panel displays a volcano plot of log₂ fold change versus –log₁₀(FDR-adjusted p-value) for all detected metabolites in the indicated treatment group: BBF | 24h, BBF | 96h, BaP | 24h, and BaP | 96h. Metabolites with a FDR-adjusted p-value < 0.05 and an absolute log₂ fold change > 1 are highlighted and annotated. Significant metabolites with increased abundance are shown in red; those with decreased abundance are shown in blue. For these metabolites, we further evaluated consistency across exposure conditions by examining the directionality of changes relative to the respective control. Pairwise group comparisons were assessed using post hoc testing, and only metabolites that demonstrated consistent directional shifts across both exposure levels- or a significant change at the high concentration that was accompanied by a similar but nonsignificant trend at the low concentration were retained for interpretation. These selected metabolites are listed in Table 3.

**Table 3.** Curated list of metabolites exhibiting significant and consistent changes in abundance across multiple concentration levels or exposure conditions. Each entry indicates the metabolite name, associated PAH (BaP or BBF), concentration, exposure time, and direction of change relative to control. Metabolites marked with an asterisk (*) and highlighted were identified as robust responders, showing consistent directional regulation (e.g., upregulation or downregulation) at high concentration, multiple concentrations, or in response to both chemicals.

| Metabolite | PAH | Concentration | Time | Direction |
| --- | --- | --- | --- | --- |
| *2-aminoadipic acid | BBF | 10 μM | 96h | downregulated |
| *2-aminoadipic acid | BBF | 1 μM | 96h | downregulated |
| acetylcarnitine (C2) | BBF | 10 μM | 96h | upregulated |
| adenine | BaP | 10 μM | 96h | downregulated |
| cystathionine | BBF | 10 μM | 96h | downregulated |
| gamma-butyrobetaine/<br>deoxycarnitine | BaP | 10 μM | 96h | upregulated |
| glutamylthreonine | BaP | 20 μM | 24h | upregulated |
| glutamylthreonine | BaP | 2 μM | 24h | upregulated |
| *glycerophosphorylinositol | BBF | 10 μM | 96h | downregulated |
| *glycerophosphorylinositol | BBF | 1 μM | 96h | downregulated |
| hexanoylcarnitine (C6) | BBF | 10 μM | 96h | downregulated |
| N-acetylglutamate | BaP | 10 μM | 96h | upregulated |
| N-acetylputrescine | BaP | 10 μM | 96h | upregulated |
| *N-acetylspermidine | BBF | 10 μM | 96h | upregulated |
| *N-acetylspermidine | BBF | 1 μM | 96h | upregulated |
| *N-acetylspermidine | BaP | 10 μM | 96h | upregulated |
| *orotidine | BBF | 10 μM | 96h | downregulated |
| *orotidine | BBF | 1 μM | 96h | downregulated |
| *S-adenosylmethionine | BBF | 10 μM | 96h | downregulated |
| *S-adenosylmethionine | BBF | 1 μM | 96h | downregulated |
| *valerylcarnitine (C5) | BBF | 10 μM | 96h | downregulated |
| *valerylcarnitine (C5) | BBF | 1 μM | 96h | downregulated |
| *beta-citrylglutamate | BBF | 10 μM | 96h | upregulated |
| *beta-citrylglutamate | BBF | 1 μM | 96h | upregulated |
| glucuronic acid | BaP | 10 μM | 96h | upregulated |

The Table 3 emphasizes metabolites exhibiting reproducible, concentration-dependent trends suggestive of potential biomarkers or mechanistic intermediates in the metabolic response to PAH exposure. These shortlisted metabolites displayed consistent and robust alterations in response to BaP and BBF exposure. Among the most reproducible responses were 2-aminoadipic acid, N-acetylspermidine, glycerophosphorylinositol, orotidine, valerylcarnitine (C5), S-adenosylmethionine, and beta-citrylglutamate each marked by directionally consistent changes across at least two BBF concentrations, and in some cases, also in BaP-exposed samples. For instance, 2-aminoadipic acid was downregulated at both 1 μM and 10 μM BBF after 96 h, indicating a consistent decrease in abundance in response to BBF exposure. Similarly, N-acetylspermidine, beta-citrylglutamate and orotidine were consistently upregulated and downregulated, respectively, across both BBF doses at 96 h, with N-acetylspermidine also upregulated in BaP-treated spheroids at 10 µM. S-adenosylmethionine and valerylcarnitine (C5) were consistently upregulated in response to BBF at both concentrations. In contrast, several metabolites showed exposure-specific or single-concentration effects and were therefore considered less robust. These included acetylcarnitine, adenine, gamma-butyrobetaine, N-acetylglutamate, N-acetylputrescine and glucuronic acid.

### 3.5 Metabolite regulation and enrichment analysis

To understand potential co-regulation among the selected metabolites (highlighted in Tables 2 and 3, as the ones being robust), we applied a network-based approach (Figure 5).

**Figure 5.**
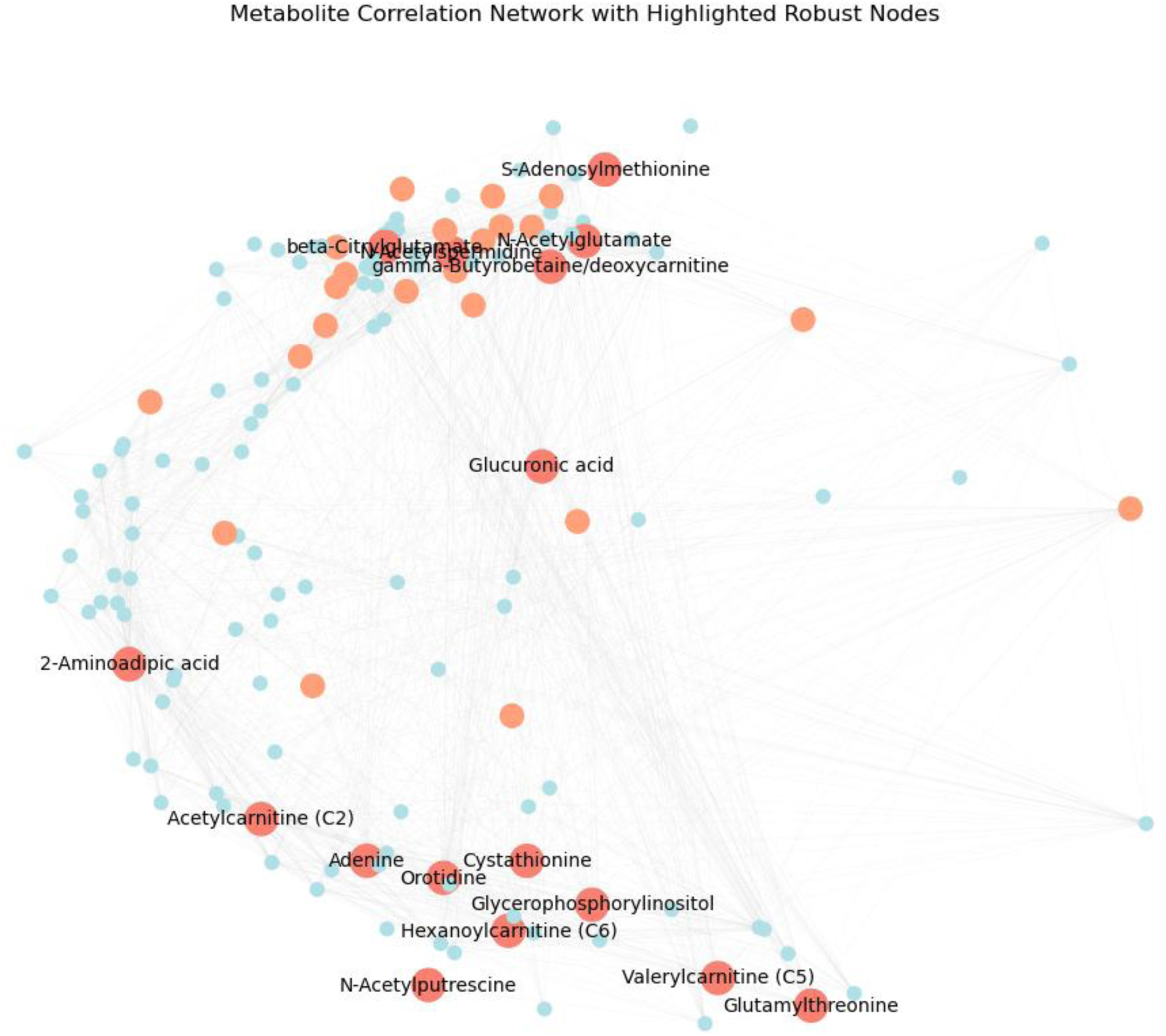
Metabolite correlation network showing community structure. Nodes represented individual metabolites, and edges represented high-confidence correlations. Metabolites highlighted in light-orange colours represent PCA-aligned, and those identified as more robust by statistical testing are shown in darker orange colour with labels.

Robustly responsive metabolites, those showing consistent, dose-dependent regulation across multiple exposure conditions, are highlighted in dark orange and labelled. These include 2-aminoadipic acid, glycerophosphorylinositol, N-acetylspermidine, orotidine, S-adenosylmethionine, valerylcarnitine (C5), beta-citrylglutamate and glucuronic acid (see Table 3).

The central or highly connected positions of these metabolites suggest potential regulatory or integrative roles in exposure-induced metabolic reprogramming. The spatial separation of these nodes across communities also hints at their involvement in multiple distinct metabolic pathways or functional modules. These results are summarized in Table 3, which served as the starting point for the enrichment analysis. From that list, we grouped the metabolites by chemical BBF or BaP exposure and ran pathway enrichment separately for each group. This allowed us to explore and compare how each PAH might affect different metabolic pathways. For the BBF group, the enrichment analysis revealed several pathways that were ranked highly based on enrichment ratio and p-values (Figure 6). Methylhistidine metabolism, for example, stood out with the strongest signal, followed by pathways such as phosphatidylcholine biosynthesis, beta oxidation of very long chain fatty acids, and spermidine and spermine biosynthesis. For the BaP group, the enrichment results were much more limited compared to BBF. We identified only two pathways from the analysis: carnitine synthesis and purine metabolism. Carnitine Synthesis had a relatively high enrichment ratio, suggesting that this pathway may be affected by BaP exposure. However, the false discovery rate values for both pathways were high, so the results were not statistically significant after correction for multiple comparisons. Purine Metabolism had a lower enrichment ratio and a higher p-value, pointing to a weaker signal. Overall, these results suggest that BaP may induce more specific or targeted changes in metabolism, unlike the broader effects observed with BBF.

**Figure 6.**
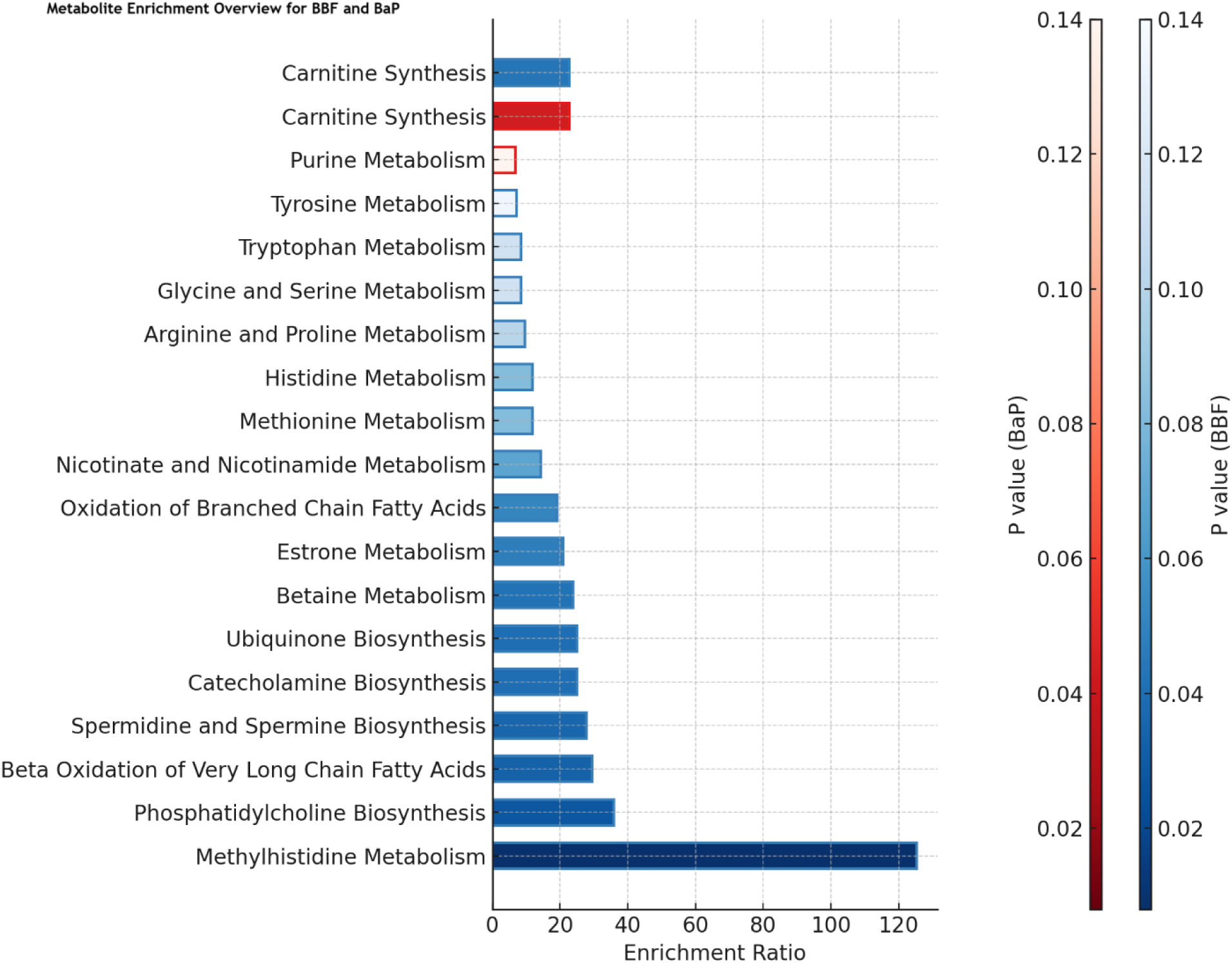
Metabolite set enrichment analysis for BaP and BBF exposed groups. The bar plot shows the most enriched metabolic pathways based on enrichment ratio and associated p-values (colour scale). In the BaP-exposed group (red), carnitine synthesis was the most significantly enriched pathway, followed by purine metabolism. In contrast, the BBF-exposed group (blue) showed broader metabolic involvement, with top-ranked pathways including methylhistidine metabolism, phosphatidylcholine biosynthesis, beta oxidation of very long chain fatty acids, and polyamine-related pathways such as spermidine and spermine biosynthesis.

While none of these passed the significance threshold after FDR correction, the raw p-values and enrichment ratios still pointed to biologically relevant changes. In the network view for the BBF group (Figure 7), most of these pathways were not scattered randomly. Instead, a large subset formed a connected cluster, suggesting that several of the affected metabolites were shared across multiple pathways. This was especially true for those involved in amino acid metabolism and methylation processes, such as methionine metabolism, betaine metabolism, histidine metabolism, and glycine and serine metabolism. This clustering is meaningful because it implies more than just isolated pathway hits; it hints at a coordinated response involving S-adenosylmethionine (SAM) and related metabolites. SAM plays a central role in methylation and polyamine biosynthesis, both of which are essential for cellular regulation and stress response.

**Figure 7.**
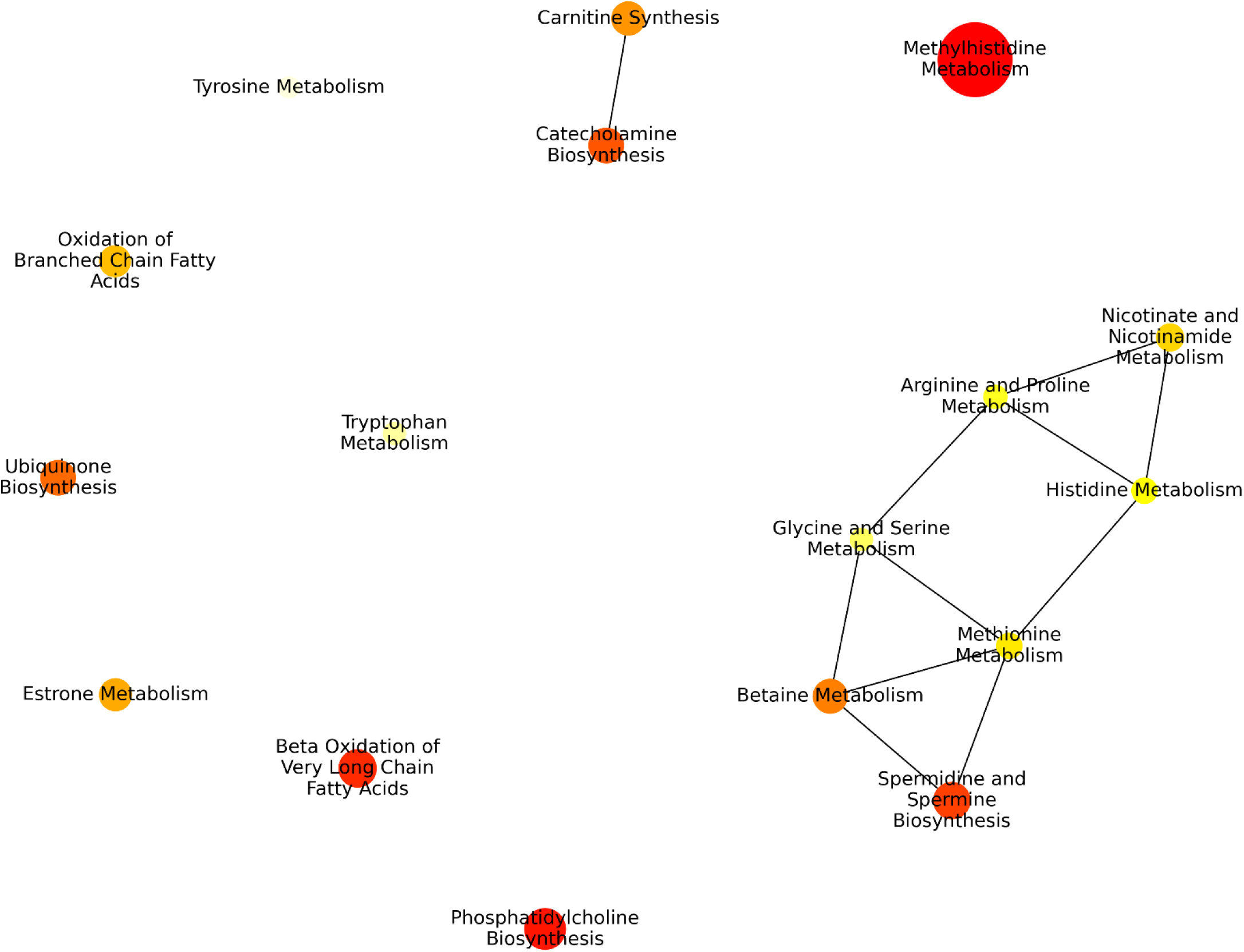
Network representation of enriched pathways in the BBF group at 96 hours. Pathways related to amino acid metabolism and methylation form a connected cluster, indicating potential biological interdependence. Figure generated in Python using metabolite set enrichment network data exported from MetaboAnalyst 6.0 (Pang et al., 2024)

Even though the statistical correction (FDR) prevents us from formally declaring these pathways significant, the biological overlap and network coherence strongly suggest that BBF may be affecting a tightly interconnected metabolic module related to methylation and amino acid metabolism.

## Discussion

Physiologically relevant *in vitro* models are increasingly essential in predictive toxicology, particularly for reducing reliance on animal testing while improving human translational relevance. A major limitation of conventional two-dimensional (2D) liver cell cultures lies in their compromised xenobiotic metabolism, which often underrepresents *in vivo* biotransformation processes (Štampar et al., 2021). To address this, we utilized a dynamic three-dimensional (3D) HepG2 spheroid model cultured in a state-of-the-art clinostat-based CelVivo system. This advanced platform enables the formation of uniform, functional spheroids that closely replicate *in vivo* liver architecture, including improved cellular organization, tissue-level gradients, and metabolic functionality. Notably, culturing HepG2 cells in this 3D environment promotes dynamic transcriptomic and metabolic maturation over time (Kim et al., 2024), leading to enhanced xenobiotic metabolism compared to standard 2D monolayers. This is particularly critical when assessing compounds such as polycyclic aromatic hydrocarbons (PAHs), whose genotoxicity relies on enzymatic bioactivation into reactive intermediates. Notably, the CelVivo system also supports long-term cultivation under physiologically relevant conditions, allowing extended exposure to low concentrations of toxicants, an essential feature for detecting subtle or delayed biological responses and for mimicking realistic human exposure scenarios (Wrzesinski et al., 2021). Together, these properties make the 3D spheroid model highly suitable for mechanistic toxicology studies of environmental contaminants with complex biotransformation profiles, such as polycyclic aromatic hydrocarbons (PAHs) (Štampar et al., 2021). Building on this physiological relevance, we performed untargeted metabolomic profiling of HepG2 spheroids exposed to two PAHs, benzo[a]pyrene (BaP) and benzo[b]fluoranthene (BBF) at both early (24 h) and prolonged (96 h) time points to unravel the cellular mechanisms underlying their toxicity. Following 24 hours of exposure, no metabolites exhibited significant alterations or emerged as robust indicators of response to either BaP or BBF. This suggests that short-term exposure may be insufficient to trigger detectable metabolic reprogramming in HepG2 spheroids. In contrast, after 96 hours of exposure, seven metabolites, namely 2-aminoadipic acid, N-acetylspermidine, glycerophosphorylinositol, orotidine, valerylcarnitine (C5), S-adenosylmethionine, and beta-citrylglutamate showed consistent and concentration-dependent changes at 96 hours in response to both BaP and BBF. Each of these metabolites displayed a uniform increase or decrease across all tested concentrations, suggesting their involvement in a lasting or delayed response to PAH exposure. These metabolites were selected for closer investigation not only because of their reproducibility, but also due to their biological relevance in cellular processes such as amino acid catabolism, lipid turnover, epigenetic regulation, cellular stress adaptation, as well as structural factors like membrane integrity. Principal component analysis (Figure 2) further supported these findings, with cells exposed to higher concentrations of BaP and BBF forming distinct clusters that were separated from the controls and low-concentration groups. By highlighting these precise and persistent metabolic shifts, our study provides a deeper understanding of how metabolically activated polycyclic aromatic hydrocarbons (PAHs) disrupt normal cellular function while simultaneously triggering compensatory responses aimed at preserving homeostasis. The contribution of these metabolites to the observed clustering highlights their potential as sensitive biomarkers of prolonged PAH-induced cellular stress. Through the characterization of these specific metabolic alterations, our study offers insight into the molecular-level disruptions caused by metabolically activated environmental toxicants. According to our results (Table 3), N-acetylspermidine levels were consistently increased in HepG2 spheroids exposed to both BaP and BBF at 1 μM and 10 μM for 96 hours. N-acetylspermidine is an intermediate of the polyamine pathway, which plays a critical role in the regulation of cell growth, differentiation, and cell death (Schibalski et al., 2024). Alterations in polyamine metabolism have been associated with cellular responses to various forms of stress and exposure to toxic compounds, including polycyclic aromatic hydrocarbons (PAHs). For example, Potratz et al., (2016) showed that PAH exposure causes oxidative stress and programmed cell death in human keratinocytes, along with changes in polyamine levels. Similarly, Masutin et al., (2022) reported that exposure to xenobiotics disrupts polyamine metabolism in skin cells, showing that the pathway is sensitive to environmental toxicants. A related observation was made by Alhonen et al., (2000) in a transgenic rat model with increased levels of the enzyme SSAT (spermidine/spermine N¹-acetyltransferase), which initiates polyamine breakdown. When the normal polyamine oxidation pathway was chemically blocked in these rats, N-acetylspermidine accumulated in the pancreas. Although this model involved genetic changes and enzyme inhibition rather than toxicant exposure, the increase in this same metabolite supports the idea that disrupted polyamine metabolism can result in cellular stress. In line with these findings, the accumulation of N-acetylspermidine in our 3D hepatic model may reflect impaired polyamine turnover, pointing to a possible stress response and loss of cellular balance following BaP and BBF exposure. Following the dysregulation observed in polyamine metabolism, we found that glycerophosphorylinositol was significantly downregulated in response to BBF at both tested concentrations after 96 h Glycerophosphorylinositol is a degradation product of inositol-containing lipids, such as phosphatidylinositol, or from glycosylphosphatidylinositol (GPI) anchors. GPI anchors attach proteins to the cell surface and support various cell functions, including cell membrane biosynthesis and cell signalling (Komath et al., 2022). PAH exposure has been widely reported to disrupt lipid metabolism, including both phospholipid and sphingolipid pathways, potentially impairing membrane integrity and signalling cascades (Smith et al., 2019; Wang et al., 2015). Our finding of decreased glycerophosphorylinositol suggests a stress-induced remodeling or depletion of membrane lipids in hepatocyte spheroids exposed to BBF, which may reflect impaired biosynthesis or increased degradation of GPI-anchored proteins. Altered turnover of these GPI-anchored proteins may contribute to the metabolic disturbances observed (Satpute-Krishnan et al., 2014). In our study, valerylcarnitine (C5) was downregulated at both 1 μM and 10 μM BBF after 96 hours of exposure. Valerylcarnitine is a short-chain acylcarnitine formed during branched-chain amino acid catabolism and plays a crucial role in transporting fatty acids into mitochondria for β-oxidation, a key energy-producing process (Koves et al., 2008). Disruptions in carnitine metabolism and β-oxidation have been reported in response to PAH exposure such as chrysene, benzo[*a*]pyrene and dibenzo[*a,l*]pyrene, which can impair mitochondrial fatty acid catabolism and contribute to energy deficits (Potratz et al., 2016; Wang et al., 2015). Similar reductions have been reported in response to environmental pollutants like in a study on short-term microplastic exposure in diabetic rats significantly decreased valerylcarnitine, correlating with immune marker changes and highlighting its sensitivity to metabolic disruption under toxic stress (Li et al., 2025). Mednova et al., (2022) observed reduced C5 levels in schizophrenia patients without metabolic syndrome, linking this to diminished mitochondrial β-oxidation capacity and broader bioenergetic imbalance. These findings support the presumption that BaP and BBF exposure interfere with fatty acid transport and mitochondrial β-oxidation in HepG2 spheroids, contributing to systemic metabolic alterations. We found that β-citryl-L-glutamate (β-CG) was significantly upregulated in HepG2 spheroids following exposure to BBF at both 1 µM and 10 µM for 96 h. β-CG is a non–proteinogenic glutamate derivative, which couples citric acid and glutamate. It functions as an endogenous iron carrier, capable of binding Fe (II) and delivering it to mitochondrial aconitase, thereby reactivating the enzyme under oxidative stress conditions (Hamada-Kanazawa et al., 2011). The observed increase in β-CG following BBF exposure may represent a cellular adaptive response aimed at preserving mitochondrial function and mitigating BBF-induced oxidative damage, as mitochondrial aconitase is particularly sensitive to reactive oxygen species (ROS) (Mansilla et al., 2023) generated by PAHs like BBF. Although data on β-CG in toxicological contexts are limited, increased β-CG levels have also been reported in metabolomic studies in muscle tissue under adaptive or stress-inducing conditions, such as resistance exercise (Gehlert et al., 2022). Taken together, our findings suggest that BBF causes an increase in β-CG as part of a mitochondrial defence mechanism, maintaining energy production during oxidative stress. We observed a significant downregulation of S-adenosylmethionine (SAM) in BBF-exposed spheroids at both concentrations after 96 hours, indicating alterations in methionine metabolism and methylation activity. SAM is the primary methyl group donor involved in various cellular processes, including the methylation of DNA, RNA, proteins, and lipids. These methylation reactions regulate gene expression, epigenetic modifications, and overall cellular homeostasis (Caracausi et al., 2024). Synthesized from methionine and ATP, SAM serves as a central molecule linking methylation to other vital pathways, such as polyamine biosynthesis and antioxidant defence (Rudenko et al., 2025). Disruptions in methionine metabolism and SAM levels have been reported in response to PAH exposure, frequently associated with oxidative stress and impaired methylation capacity (Gao et al., 2024; Ma et al., 2024). Supporting this, previous studies have demonstrated that environmental pollutants such as PAHs can disrupt the methionine cycle, notably by reducing the expression of methionine adenosyltransferase 1A (MAT1A), thereby impairing SAM synthesis and subsequent methylation reactions (Rider and Carlsten, 2019). Interestingly, although our study included BaP-treated spheroids, we did not observe a significant downregulation of SAM in these samples. However, a study by Wang et al., (2023) describes a mechanism whereby BaP increases glycine N-methyltransferase (GNMT) activity, enhancing the conversion of SAM to S-adenosylhomocysteine (SAH), thereby reducing SAM availability and ultimately leading to global DNA hypomethylation. This divergence suggests that the effect of BaP on SAM levels may be context-dependent, potentially influenced by factors such as exposure duration, concentration, cell type, or model system. Orotidine levels were significantly downregulated in HepG2 spheroids exposed to both 1 μM and 10 μM BBF for 96 hours, indicating a shift in nucleotide metabolism, specifically in the pyrimidine biosynthesis pathway. Orotidine is an intermediate in the *de novo* synthesis of pyrimidine nucleotides, which are essential for RNA and DNA synthesis (Yadav et al., 2020). Polycyclic aromatic hydrocarbons such as BBF are known to generate reactive oxygen species (ROS) (Sombiri et al., 2024), which can damage mitochondrial function and inhibit key enzymes involved in pyrimidine biosynthesis, including dihydroorotate dehydrogenase (DHODH), a mitochondrial enzyme critical for converting dihydroorotate to orotate, a precursor of orotidine (Zhou et al., 2021). Dysfunction of DHODH leads to decreased orotate and subsequently orotidine levels, disrupting nucleotide synthesis and energy metabolism. Furthermore, oxidative stress and DNA damage induced by BBF activate cell cycle checkpoints that reduce cellular proliferation and nucleotide demand, contributing to decreased flux through the de novo pyrimidine pathway (Hubackova et al., 2020). Cells under such stress often compensate by upregulating salvage pathways to recycle nucleotides, which may further reduce the accumulation of intermediates like orotidine (Walter and Herr, 2022). Collectively, these mechanisms suggest that orotidine downregulation in BBF-treated HepG2 spheroids is a multifactorial consequence of mitochondrial dysfunction, oxidative stress, DNA damage responses, and metabolic reprogramming, positioning orotidine as a potential biomarker of PAH-induced hepatotoxicity. Another robustly altered metabolite identified in our analysis was 2-aminoadipic acid (2-AAA), which was significantly downregulated in HepG2 spheroids exposed to BBF at both 1 μM and 10 μM concentrations for 96 h. 2-AAA is a non-proteinogenic amino acid formed during lysine catabolism via the saccharopine pathway. It is synthesized by the enzyme aminoadipate-semialdehyde synthase (AASS) and subsequently converted into α-ketoadipate, which enters the tricarboxylic acid (TCA) cycle, linking lysine catabolism to energy metabolism (Leandro et al., 2022). The observed downregulation of 2-AAA may indicate suppression of lysine degradation or impaired flux into the TCA cycle, possibly reflecting mitochondrial dysfunction induced by BBF (Schneider et al., 2022). Since mitochondrial oxidative phosphorylation is sensitive to PAH-induced reactive oxygen species (ROS), disruption of amino acid catabolism could contribute to broader energy deficits (Murphy, 2009). Furthermore, lysine-derived metabolites like 2-AAA have been implicated in epigenetic regulation and redox balance, suggesting that their depletion may exacerbate oxidative stress or interfere with histone modification processes (Gavin and Sharma, 2010). Potratz et al., (2016) also reported a decrease in α-aminoadipic acid, which is consistent with our findings. Importantly, our metabolomic profiling further revealed changes in methylation-related metabolites and nucleotide metabolism, suggesting broader epigenetic and genomic effects of PAH exposure. While our study provides valuable insights into the metabolic alterations induced by BaP and BBF exposure, several limitations should be acknowledged. Although non-targeted metabolomics enables the detection of a wide array of metabolites across diverse pathways, it may miss specific metabolites, particularly those in low abundance or those outside the analytical sensitivity range. As a result, Specific pathways or metabolite classis may have been unintentionally underrepresented due to methodological constraints, such as limitations of the selected analytical method. Furthermore, our study was limited to two exposure durations (24 hours and 96 hours) and fixed concentrations. These conditions, while informative, may not fully capture the complexity of real-world human exposures, which are often chronic, low-dose, and intermittent. The use of HepG2 spheroids, while advantageous due to their metabolic activity and structural relevance, still represents the use of an immortalized cancer cell line. Consequently, differences in metabolic responses compared to primary human hepatocytes or other liver cell types may limit the broader applicability of the findings. Another consideration is the exclusion of specific metabolites during correlation analysis. While this step was necessary to enhance statistical robustness and reduce false positives, it may have inadvertently removed biologically relevant metabolites, potentially impacting pathway enrichment results. The conservative statistical approach, although rigorous, might have obscured subtle yet meaningful signals. Future research could expand on these findings by examining long-term, low-dose exposures that more closely mimic real-world environmental and occupational settings. Moreover, the consistent and dominant metabolic impact of BBF across concentrations highlights the need for further mechanistic studies and comparative analyses with other PAHs. These findings also highlight the potential of integrating metabolic biomarkers into Integrated Lifetime Cancer Risk (ILCR) assessments (Nisbet and LaGoy, 1992), supporting their use in regulatory toxicology and public health decision-making.

## Conclusion

Our findings demonstrate that human hepatic HepG2 spheroids represent a robust and metabolically competent experimental model for investigating the effects of airborne polycyclic aromatic hydrocarbons (PAHs). By integrating high-resolution untargeted metabolomics with multivariate statistical analysis, we uncovered distinct yet partially overlapping metabolic signatures for benzo[a]pyrene (BaP) and benzo[b]fluoranthene (BBF), identifying both compound-specific and shared metabolic responses, involving lipid metabolism, nucleotide biosynthesis, and redox balance. Principal component analysis and complementary network analysis revealed distinct clustering patterns between BaP- and BBF-exposed spheroids, indicating compound-specific metabolic responses. Notably, BBF exposure elicited a broader and more pronounced spectrum of metabolic perturbations compared to BaP, even though BaP is widely regarded as one of the most potent and representative PAHs in toxicological research and regulatory frameworks. Crucially, these differences only became evident after prolonged exposure (96 h), highlighting the value of extended-duration assays for capturing delayed or cumulative metabolic stress. Short-term exposures may therefore underestimate the risk posed by the delayed impact of specific PAHs like BBF. Complementary network analysis further supported functional divergence, highlighting functional alterations in key pathways related to lipid metabolism, nucleotide biosynthesis, and redox homeostasis. Although BBF induced broader metabolic perturbations, a subset of metabolites and pathways was commonly affected by both BaP and BBF. This overlap suggests that, despite differences in the extent and intensity of their effects, the two PAHs share underlying mechanisms of toxicity or convergent metabolic stress responses. These findings emphasize the value of integrating unsupervised statistical modelling with biologically relevant *in vitro* 3D cell systems to unravel both compound-specific and shared cellular responses to environmental toxicants. Our study not only highlights the use of spheroids for mechanistic toxicological studies but also provides a framework for identifying potential metabolic biomarkers of PAH exposure. Future research should focus on validating these findings under exposure scenarios that more closely mimic human environmental and occupational conditions, thereby improving their applicability for risk assessment and regulatory decision-making concerning airborne toxicants. Importantly, the metabolic perturbations observed, such as the upregulation of 2-aminoadipic acid, reflect metabolic changes commonly associated with metabolic disorders like metabolic syndrome and type 2 diabetes. These results deepen our understanding of how environmental pollutants may contribute to widespread metabolic dysregulation in humans, offering valuable avenues for preventive strategies and more accurate approaches to human health risk assessment. Together, these findings not only advance toxicological research but also highlight the urgency of addressing environmental pollutant exposure as a contributor to global metabolic health challenges.

## Supplementary Information

The supplementary dataset for this study can be accessed on Zenodo: Bošnjaković, A. (2025). *Supplementary Excel Table for “Metabolomic fingerprints of PAH exposure: Identifying toxicological biomarkers in dynamically cultured 3D cell spheroids”*. Zenodo. https://doi.org/10.5281/zenodo.15851269

## Acknowledgements

The experimental work was funded by Horizon Europe (EDIAQI project #101057497; CutCancer project #10050740). ML has received funding from the Croatian Science Foundation (MOBODL-2023-12-5502). AB has received funding from the Croatian Science Foundation (MOBDOK-2023-1077. BZ received funding from the Slovenian Research Agency (P1-0245).

## References

Alhonen, L., Parkkinen, J.J., Keinänen, T., Sinervirta, R., Herzig, K.-H., Jänne, J., 2000. Activation of polyamine catabolism in transgenic rats induces acute pancreatitis. Proc. Natl. Acad. Sci. 97, 8290–8295. 10.1073/pnas.140122097

Authority (EFSA), E.F.S., 2008. Polycyclic Aromatic Hydrocarbons in Food - Scientific Opinion of the Panel on Contaminants in the Food Chain. EFSA J. 6, 724. 10.2903/j.efsa.2008.724

Bai, H., Wu, M., Zhang, H., Tang, G., 2017. Chronic polycyclic aromatic hydrocarbon exposure causes DNA damage and genomic instability in lung epithelial cells. Oncotarget 8, 79034– 79045. 10.18632/oncotarget.20891

Bao, Q., Zhao, S., Liu, Y., Yao, Y., You, J., Xiong, J., 2023. Benzo[b]fluoranthene induced oxidative stress and apoptosis in human airway epithelial cells via mitochondrial disruption. J. Appl. Toxicol. 43, 1083–1094. 10.1002/jat.4445

Caracausi, M., Ramacieri, G., Catapano, F., Cicilloni, M., Lajin, B., Pelleri, M.C., Piovesan, A., Vitale, L., Locatelli, C., Pirazzoli, G.L., Strippoli, P., Antonaros, F., Vione, B., 2024. The functional roles of S-adenosyl-methionine and S-adenosyl-homocysteine and their involvement in trisomy 21. BioFactors 50, 709–724. 10.1002/biof.2044

Chen, C.-H.S., Yuan, T.-H., Shie, R.-H., Wu, K.-Y., Chan, C.-C., 2017. Linking sources to early effects by profiling urine metabolome of residents living near oil refineries and coal-fired power plants. Environ. Int. 102, 87–96. 10.1016/j.envint.2017.02.003

Choi, H., Harrison, R., Komulainen, H., Saborit, J.M.D., 2010. Polycyclic aromatic hydrocarbons, in: WHO Guidelines for Indoor Air Quality: Selected Pollutants. World Health Organization.

Clauset, A., Newman, M.E.J., Moore, C., 2004. Finding community structure in very large networks. Phys. Rev. E 70, 066111. 10.1103/PhysRevE.70.066111

Deng, Z., Li, L., Jing, Z., Luo, X., Yu, F., Zeng, W., Bi, W., Zou, J., 2024. Association between environmental phthalates exposure and gut microbiota and metabolome in dementia with Lewy bodies. Environ. Int. 190, 108806. 10.1016/j.envint.2024.108806

Ewa, B., Danuta, M.-Š., 2017. Polycyclic aromatic hydrocarbons and PAH-related DNA adducts. J. Appl. Genet. 58, 321–330. 10.1007/s13353-016-0380-3

Fang, Y., Yin, W., He, C., Shen, Q., Xu, Y., Liu, C., Zhou, Y., Liu, G., Zhao, Y., Zhang, H., Zhao, K., 2024. Adverse impact of phthalate and polycyclic aromatic hydrocarbon mixtures on birth outcomes: A metabolome Exposome-Wide association study. Environ. Pollut. Barking Essex 1987 357, 124460. 10.1016/j.envpol.2024.124460

Fey, S.J., Wrzesinski, K., 2012. Determination of Drug Toxicity Using 3D Spheroids Constructed From an Immortal Human Hepatocyte Cell Line. Toxicol. Sci. 127, 403–411. 10.1093/toxsci/kfs122

Gajski, G., Gerić, M., Žegura, B., Novak, M., Nunić, J., Bajrektarević, D., Garaj-Vrhovac, V., Filipič, M., 2016. Genotoxic potential of selected cytostatic drugs in human and zebrafish cells. Environ. Sci. Pollut. Res. 23, 14739–14750. 10.1007/s11356-015-4592-6

Gao, P., da Silva, E., Hou, L., Denslow, N.D., Xiang, P., Ma, L.Q., 2018. Human exposure to polycyclic aromatic hydrocarbons: Metabolomics perspective. Environ. Int. 119, 466–477. 10.1016/j.envint.2018.07.017

Gao, R., Jiang, Z., Wu, X., Cai, Z., Sang, N., 2024. Metabolic regulation of tumor cells exposed to different oxygenated polycyclic aromatic hydrocarbons. Sci. Total Environ. 907, 167833. 10.1016/j.scitotenv.2023.167833

Gavin, D.P., Sharma, R.P., 2010. Histone modifications, DNA methylation, and Schizophrenia. Neurosci. Biobehav. Rev., Special Section: Developmental determinants of sensitivity and resistance to stress: A tribute to Seymour “Gig” Levine 34, 882–888. 10.1016/j.neubiorev.2009.10.010

Gehlert, S., Weinisch, P., Römisch-Margl, W., Jaspers, R.T., Artati, A., Adamski, J., Dyar, K.A., Aussieker, T., Jacko, D., Bloch, W., Wackerhage, H., Kastenmüller, G., 2022. Effects of Acute and Chronic Resistance Exercise on the Skeletal Muscle Metabolome. Metabolites 12, 445. 10.3390/metabo12050445

Guo, P., Furnary, T., Vasiliou, V., Yan, Q., Nyhan, K., Jones, D.P., Johnson, C.H., Liew, Z., 2022. Non-targeted metabolomics and associations with per- and polyfluoroalkyl substances (PFAS) exposure in humans: A scoping review. Environ. Int. 162, 107159. 10.1016/j.envint.2022.107159

Hagberg, A.A., Schult, D.A., Swart, P.J., 2008. Exploring Network Structure, Dynamics, and Function using NetworkX. Presented at the Python in Science Conference, Pasadena, California, pp. 11–15. 10.25080/TCWV9851

Hamada-Kanazawa, M., Narahara, M., Takano, M., Min, K.S., Tanaka, K., Miyake, M., 2011. β-Citryl-L-glutamate acts as an iron carrier to activate aconitase activity. Biol. Pharm. Bull. 34, 1455–1464. 10.1248/bpb.34.1455

Hubackova, S., Davidova, E., Boukalova, S., Kovarova, J., Bajzikova, M., Coelho, A., Terp, M.G., Ditzel, H.J., Rohlena, J., Neuzil, J., 2020. Replication and ribosomal stress induced by targeting pyrimidine synthesis and cellular checkpoints suppress p53-deficient tumors. Cell Death Dis. 11, 110. 10.1038/s41419-020-2224-7

IARC, n.d. IARC Monographs on the Identification of Carcinogenic Hazards to Humans [WWW Document]. URL https://monographs.iarc.who.int/list-of-classifications (accessed 4.17.25).

India-Aldana, S., Yao, M., Midya, V., Colicino, E., Chatzi, L., Chu, J., Gennings, C., Jones, D.P., Loos, R.J.F., Setiawan, V.W., Smith, M.R., Walker, R.W., Barupal, D., Walker, D.I., Valvi, D., 2023. PFAS Exposures and the Human Metabolome: A Systematic Review of Epidemiological Studies. Curr. Pollut. Rep. 9, 510–568. 10.1007/s40726-023-00269-4

Jakovljević, I., Sever Štrukil, Z., Godec, R., Bešlić, I., Davila, S., Lovrić, M., Pehnec, G., 2020. Pollution Sources and Carcinogenic Risk of PAHs in PM1 Particle Fraction in an Urban Area. Int. J. Environ. Res. Public. Health 17, 9587. 10.3390/ijerph17249587

Jakovljević, I., Štrukil, Z.S., Pehnec, G., Horvat, T., Sanković, M., Šumanovac, A., Davila, S., Račić, N., Gajski, G., 2025. Ambient air pollution and carcinogenic activity at three different urban locations. Ecotoxicol. Environ. Saf. 289, 117704. 10.1016/j.ecoenv.2025.117704

Kazensky, L., Matković, K., Gerić, M., Žegura, B., Pehnec, G., Gajski, G., 2024. Impact of indoor air pollution on DNA damage and chromosome stability: a systematic review. Arch. Toxicol. 10.1007/s00204-024-03785-4

Kim, C., Zhu, Z., Barbazuk, W.B., Bacher, R.L., Vulpe, C.D., 2024. Time-course characterization of whole-transcriptome dynamics of HepG2/C3A spheroids and its toxicological implications. Toxicol. Lett. 401, 125–138. 10.1016/j.toxlet.2024.10.004

Komath, S.S., Fujita, M., Hart, G.W., Ferguson, M.A.J., Kinoshita, T., 2022. Glycosylphosphatidylinositol Anchors, in: Varki, A., Cummings, R.D., Esko, J.D., Stanley, P., Hart, G.W., Aebi, M., Mohnen, D., Kinoshita, T., Packer, N.H., Prestegard, J.H., Schnaar, R.L., Seeberger, P.H. (Eds.), Essentials of Glycobiology. Cold Spring Harbor Laboratory Press, Cold Spring Harbor (NY).

Koves, T.R., Ussher, J.R., Noland, R.C., Slentz, D., Mosedale, M., Ilkayeva, O., Bain, J., Stevens, R., Dyck, J.R.B., Newgard, C.B., Lopaschuk, G.D., Muoio, D.M., 2008. Mitochondrial Overload and Incomplete Fatty Acid Oxidation Contribute to Skeletal Muscle Insulin Resistance. Cell Metab. 7, 45–56. 10.1016/j.cmet.2007.10.013

Leandro, J., Khamrui, S., Suebsuwong, C., Chen, P.-J., Secor, C., Dodatko, T., Yu, C., Sanchez, R., DeVita, R.J., Houten, S.M., Lazarus, M.B., 2022. Characterization and structure of the human lysine-2-oxoglutarate reductase domain, a novel therapeutic target for treatment of glutaric aciduria type 1. 10.1101/2022.05.20.492856

Li, M., Ye, G., Liu, Y., Yang, T., Zhao, B., Jiang, R., Chen, G., 2025. Short-term microplastic exposure: A double whammy to lung metabolism and fecal microflora in diabetic SD rats. Ecotoxicol. Environ. Saf. 297, 118229. 10.1016/j.ecoenv.2025.118229

Lovrić, M., Gajski, G., Fernández-Agüera, J., Pöhlker, M., Gursch, H., Consortium, T.E., Borg, A., Switters, J., Mureddu, F., 2024a. Evidence-driven indoor air quality improvement: An innovative and interdisciplinary approach to improving indoor air quality. BioFactors n/a. 10.1002/biof.2126

Lovrić, M., Horner, D., Chen, L., Brustad, N., Malby Schoos, A.-M., Lasky-Su, J., Chawes, B., Rasmussen, M.A., 2024b. Vertical Metabolome Transfer from Mother to Child: An Explainable Machine Learning Method for Detecting Metabolomic Heritability. Metabolites 14, 136. 10.3390/metabo14030136

Lovrić, M., Račić, N., Pehnec, G., Horvat, T., Lovrić Štefiček, M.J., Jakovljević, I., 2024c. Indoor Polycyclic Aromatic Hydrocarbons—Relationship to Ambient Air, Risk Estimation, and Source Apportionment Based on Household Measurements. Atmosphere 15, 1525. 10.3390/atmos15121525

Ma, J., Yu, H., Li, G., An, T., 2024. Mechanism of cytochrome P450s mediated interference with glutathione and amino acid metabolisms from halogenated PAHs exposure. J. Hazard. Mater. 473, 134589. 10.1016/j.jhazmat.2024.134589

Mallah, Manthar Ali, Changxing, L., Mallah, Mukhtiar Ali, Noreen, S., Liu, Y., Saeed, M., Xi, H., Ahmed, B., Feng, F., Mirjat, A.A., Wang, W., Jabar, A., Naveed, M., Li, J.-H., Zhang, Q., 2022. Polycyclic aromatic hydrocarbon and its effects on human health: An overeview. Chemosphere 296, 133948. 10.1016/j.chemosphere.2022.133948

Mallah, Manthar Ali, Mallah, Mukhtiar Ali, Liu, Y., Xi, H., Wang, W., Feng, F., Zhang, Q., 2021. Relationship Between Polycyclic Aromatic Hydrocarbons and Cardiovascular Diseases: A Systematic Review. Front. Public Health 9, 763706. 10.3389/fpubh.2021.763706

Mansilla, S., Tórtora, V., Pignataro, F., Sastre, S., Castro, I., Chiribao, Ma.L., Robello, C., Zeida, A., Santos, J., Castro, L., 2023. Redox sensitive human mitochondrial aconitase and its interaction with frataxin: *In vitro* and *in silico* studies confirm that it takes two to tango. Free Radic. Biol. Med. 197, 71–84. 10.1016/j.freeradbiomed.2023.01.028

Masutin, V., Kersch, C., Schmitz-Spanke, S., 2022. A systematic review: metabolomics-based identification of altered metabolites and pathways in the skin caused by internal and external factors. Exp. Dermatol. 31, 700–714. 10.1111/exd.14529

Mednova, I.A., Chernonosov, A.A., Kornetova, E.G., Semke, A.V., Bokhan, N.A., Koval, V.V., Ivanova, S.A., 2022. Levels of Acylcarnitines and Branched-Chain Amino Acids in Antipsychotic-Treated Patients with Paranoid Schizophrenia with Metabolic Syndrome. Metabolites 12, 850. 10.3390/metabo12090850

MetaboAnalyst 6.0: towards a unified platform for metabolomics data processing, analysis and interpretation | Nucleic Acids Research | Oxford Academic [WWW Document], n.d. URL https://academic.oup.com/nar/article/52/W1/W398/7642060 (accessed 5.26.25).

Murphy, M.P., 2009. How mitochondria produce reactive oxygen species. Biochem. J. 417, 1–13. 10.1042/BJ20081386

Nisbet, I.C.T., LaGoy, P.K., 1992. Toxic equivalency factors (TEFs) for polycyclic aromatic hydrocarbons (PAHs). Regul. Toxicol. Pharmacol. 16, 290–300. 10.1016/0273-2300(92)90009-X

Pang, Z., Lu, Y., Zhou, G., Hui, F., Xu, L., Viau, C., Spigelman, A.F., MacDonald, P.E., Wishart, D.S., Li, S., Xia, J., 2024. MetaboAnalyst 6.0: towards a unified platform for metabolomics data processing, analysis and interpretation. Nucleic Acids Res. 52, W398–W406. 10.1093/nar/gkae253

Pezdirc, M., Žegura, B., Filipič, M., 2013. Genotoxicity and induction of DNA damage responsive genes by food-borne heterocyclic aromatic amines in human hepatoma HepG2 cells. Food Chem. Toxicol. 59, 386–394. 10.1016/j.fct.2013.06.030

Piccinini, F., Tesei, A., Arienti, C., Bevilacqua, A., 2015. Cancer multicellular spheroids: Volume assessment from a single 2D projection. Comput. Methods Programs Biomed. 118, 95–106. 10.1016/j.cmpb.2014.12.003

Potratz, S., Jungnickel, H., Grabiger, S., Tarnow, P., Otto, W., Fritsche, E., von Bergen, M., Luch, A., 2016. Differential cellular metabolite alterations in HaCaT cells caused by exposure to the aryl hydrocarbon receptor-binding polycyclic aromatic hydrocarbons chrysene, benzo[*a*]pyrene and dibenzo[*a,l*]pyrene. Toxicol. Rep. 3, 763–773. 10.1016/j.toxrep.2016.09.003

Rider, C.F., Carlsten, C., 2019. Air pollution and DNA methylation: effects of exposure in humans. Clin. Epigenetics 11, 131. 10.1186/s13148-019-0713-2

Rudenko, A.Y., Mariasina, S.S., Ozhiganov, R.M., Sergiev, P.V., Polshakov, V.I., 2025. Enzymatic Reactions of S-Adenosyl-L-Methionine: Synthesis and Applications. Biochem. Biokhimiia 90, S105–S134. 10.1134/S0006297924604210

Ryu, J.Y., Hong, D.H., 2024. Association of mixed polycyclic aromatic hydrocarbons exposure with oxidative stress in Korean adults. Sci. Rep. 14, 7511. 10.1038/s41598-024-58263-9

Satpute-Krishnan, P., Ajinkya, M., Bhat, S., Itakura, E., Hegde, R.S., Lippincott-Schwartz, J., 2014. ER Stress-Induced Clearance of Misfolded GPI-Anchored Proteins via the Secretory Pathway. Cell 158, 522–533. 10.1016/j.cell.2014.06.026

Schibalski, R.S., Shulha, A.S., Tsao, B.P., Palygin, O., Ilatovskaya, D.V., 2024. The role of polyamine metabolism in cellular function and physiology. Am. J. Physiol.-Cell Physiol. 327, C341–C356. 10.1152/ajpcell.00074.2024

Schneider, J.G., Tozzo, E., Chakravarthy, M.V., 2022. Editorial: Mitochondrial Biology and Its Role in Metabolic Diseases. Front. Endocrinol. 13, 944728. 10.3389/fendo.2022.944728

Schuster, D.M., LeBlanc, D.P.M., Zhou, G., Meier, M.J., Dodge, A.E., White, P.A., Long, A.S., Williams, A., Hobbs, C., Diesing, A., Smith-Roe, S.L., Salk, J.J., Marchetti, F., Yauk, C.L., 2024. Dose-Related Mutagenic and Clastogenic Effects of Benzo[b]fluoranthene in Mouse Somatic Tissues Detected by Duplex Sequencing and the Micronucleus Assay. Environ. Sci. Technol. 58, 21450–21463. 10.1021/acs.est.4c07236

Smith, M.R., Walker, D.I., Uppal, K., Utell, M.J., Hopke, P.K., Mallon, T.M., Krahl, P.L., Rohrbeck, P., Go, Y.-M., Jones, D.P., 2019. Benzo[a]pyrene Perturbs Mitochondrial and Amino Acid Metabolism in Lung Epithelial Cells and Has Similar Correlations With Metabolic Changes in Human Serum. J. Occup. Environ. Med. 61 Suppl 12, S73–S81. 10.1097/JOM.0000000000001687

Sombiri, S., Balhara, N., Attri, D., Kharb, I., Giri, A., 2024. An overview on occurrence of polycyclic aromatic hydrocarbons in food chain with special emphasis on human health ailments. Discov. Environ. 2, 87. 10.1007/s44274-024-00121-6

Štampar, M., Sedighi Frandsen, H., Rogowska-Wrzesinska, A., Wrzesinski, K., Filipič, M., Žegura, B., 2021. Hepatocellular carcinoma (HepG2/C3A) cell-based 3D model for genotoxicity testing of chemicals. Sci. Total Environ. 755, 143255. 10.1016/j.scitotenv.2020.143255

Štampar, M., Žegura, B., 2024. In vitro hepatic 3D cell models and their application in genetic toxicology: A systematic review. Mutat. Res. Genet. Toxicol. Environ. Mutagen. 900, 503835. 10.1016/j.mrgentox.2024.503835

Topinka, J., Marvanová, S., Vondráček, J., Sevastyanova, O., Nováková, Z., Krčmář, P., Pěnčíková, K., Machala, M., 2008. DNA adducts formation and induction of apoptosis in rat liver epithelial ‘stem-like’ cells exposed to carcinogenic polycyclic aromatic hydrocarbons. Mutat. Res. Mol. Mech. Mutagen. 638, 122–132. 10.1016/j.mrfmmm.2007.09.004

US EPA, O., 2013. Guidelines for Carcinogen Risk Assessment [WWW Document]. URL https://www.epa.gov/risk/guidelines-carcinogen-risk-assessment (accessed 6.16.25).

Walter, M., Herr, P., 2022. Re-Discovery of Pyrimidine Salvage as Target in Cancer Therapy. Cells 11, 739. 10.3390/cells11040739

Wang, H., Liu, B., Chen, H., Xu, P., Xue, H., Yuan, J., 2023. Dynamic changes of DNA methylation induced by benzo(a)pyrene in cancer. Genes Environ. 45, 21. 10.1186/s41021-023-00278-1

Wang, Z., May, S.M., Charoenlap, S., Pyle, R., Ott, N.L., Mohammed, K., Joshi, A.Y., 2015. Effects of secondhand smoke exposure on asthma morbidity and health care utilization in children: a systematic review and meta-analysis. Ann. Allergy Asthma Immunol. Off. Publ. Am. Coll. Allergy Asthma Immunol. 115, 396–401.e2. 10.1016/j.anai.2015.08.005

Wojdyla, K., Wrzesinski, K., Williamson, J., Fey, S.J., Rogowska-Wrzesinska, A., 2016. Acetaminophen-induced S-nitrosylation and S-sulfenylation signalling in 3D cultured hepatocarcinoma cell spheroids. Toxicol. Res. 5, 905–920. 10.1039/c5tx00469a

Wold, S., Esbensen, K., Geladi, P., 1987. Principal component analysis. Chemom. Intell. Lab. Syst., Proceedings of the Multivariate Statistical Workshop for Geologists and Geochemists 2, 37–52. 10.1016/0169-7439(87)80084-9

Wrzesinski, K., Alnøe, S., Jochumsen, H.H., Mikkelsen, K., Bryld, T.D., Vistisen, J.S., Alnøe, P.W., Fey, S.J., Wrzesinski, K., Alnøe, S., Jochumsen, H.H., Mikkelsen, K., Bryld, T.D., Vistisen, J.S., Alnøe, P.W., Fey, S.J., 2021. A Purpose-Built System for Culturing Cells as In Vivo Mimetic 3D Structures, in: Biomechanics and Functional Tissue Engineering. IntechOpen. 10.5772/intechopen.96091

Wrzesinski, K., Fey, S.J., 2018. Metabolic Reprogramming and the Recovery of Physiological Functionality in 3D Cultures in Micro-Bioreactors. Bioengineering 5, 22. 10.3390/bioengineering5010022

Wrzesinski, K., Fey, S.J., 2013. After trypsinisation, 3D spheroids of C3A hepatocytes need 18 days to re-establish similar levels of key physiological functions to those seen in the liver. Toxicol. Res. 2, 123–135. 10.1039/c2tx20060k

Yadav, M., Kumar, R., Krishnamurthy, R., 2020. Chemistry of Abiotic Nucleotide Synthesis. Chem. Rev. 120, 4766–4805. 10.1021/acs.chemrev.9b00546

Zhou, Yue, Tao, L., Zhou, X., Zuo, Z., Gong, J., Liu, X., Zhou, Yang, Liu, C., Sang, N., Liu, H., Zou, J., Gou, K., Yang, X., Zhao, Y., 2021. DHODH and cancer: promising prospects to be explored. Cancer Metab. 9, 22. 10.1186/s40170-021-00250-z

